# An end-to-end workflow to study newly synthesized mRNA following rapid protein depletion in *Saccharomyces cerevisiae*

**DOI:** 10.1101/2024.01.26.577353

**Authors:** John B. Ridenour, Rafal Donczew

## Abstract

Accurate regulation of gene transcription by RNA polymerase II is essential for the growth and development of eukaryotic cells. Although significant progress has been made in understanding the mechanisms that regulate transcription, many questions remain unanswered. Defining the direct effects of transcriptional regulators is critically important to answering these questions. An effective approach for identifying the direct targets of transcriptional regulators is combining rapid protein depletion and quantification of newly transcribed RNA. The auxin-inducible degron (AID) system and thiol (SH)-linked alkylation for the metabolic sequencing of RNA (SLAM-seq) are powerful methods to rapidly degrade a target protein and directly quantify newly transcribed RNA, respectively. Both methods have been widely applied to study transcriptional regulation. To address unresolved questions in transcription, we engineered an end-to-end workflow in *Saccharomyces cerevisiae* to deplete proteins of interest using the AID system and measure newly transcribed RNA using SLAM-seq. We provide a step-by-step protocol to support rapid implementation and demonstrate that the workflow can help define the direct effects of transcriptional regulators using the BET proteins Bdf1 and Bdf2 as a test case. This workflow will help address outstanding questions underlying the molecular basis of transcription and other biological processes in *S. cerevisiae* and other systems.

## INTRODUCTION

Gene transcription by RNA polymerase II (Pol II) is a fundamental process in eukaryotic cells. The precise regulation of transcription is necessary for cellular growth and development and requires the coordinated activity of numerous proteins and protein complexes (1, 2). For example, Pol II alone cannot initiate transcription but must cooperate with a large set of general transcription factors (GTFs), including TFIIA, TFIIB, TFIID, TFIIE, TFIIF, and TFIIH, to perform this essential function (2, 3). In addition to basal factors such as GTFs, sequence-specific transcription factors, transcriptional coactivators, and chromatin modifiers have critical roles in regulating transcription in response to developmental or environmental cues (1, 4–6). Despite considerable progress in understanding the molecular basis of transcriptional regulation, many questions remain unanswered. The budding yeast *Saccharomyces cerevisiae* remains an excellent system for addressing unresolved questions in transcription, as the underlying biology is broadly conserved across eukaryotes.

Defining the direct effects of transcriptional regulators is critical for elucidating their function (7, 8). The rapid and specific degradation of target proteins is a powerful approach for identifying the direct targets of transcriptional regulators. The auxin-inducible degron (AID) system is one method for rapid protein degradation widely used to study transcriptional regulation in eukaryotes, including *S. cerevisiae* (8–11). Derived from a plant-specific module of the conserved eukaryotic SKP1-CUL1-F-box (SCF) E3 ubiquitin ligase complex, the AID system requires the expression of the plant F-box protein TIR1 and the genetic fusion of an AID tag to a target protein to function outside of plants (11). The addition of the plant hormone auxin promotes the association of TIR1 with the AID tag, recruitment of endogenous SCF complex components, and degradation of the target protein via the 26S proteasome. Unlike most classical genetic approaches, essential genes or synthetic lethal interactions can be studied using the AID system. We and others have used the AID system to achieve specific degradation of chromatin readers, chromatin remodeling factors, transcription factors, and transcriptional coactivators in *S. cerevisiae* (9, 12–15).

RNA synthesized immediately following a perturbation, such as rapid protein degradation, more accurately reflects the direct effects of the target factor than steady-state RNA (7, 8). Several methods selectively quantify nascent or newly synthesized RNA, including GRO-seq, PRO-seq, NET-seq, TT-seq, TimeLapse-seq, and 4sU-seq (and the equivalent method 4tU-seq) (16, 17). We and others have used 4sU-seq and 4tU-seq to quantify newly synthesized RNA following targeted protein depletion in *S. cerevisiae* (9, 12, 18–20). These methods involve labeling newly synthesized RNA with 4-thiouridine (4sU) or 4-thiouracil (4tU), followed by biochemical purification of labeled RNA (9, 21, 22). Thiol (<u>S</u>H)-linked <u>a</u>lkylation for the <u>m</u>etabolic sequencing of RNA (SLAM-seq) is an alternative method that directly quantifies 4sU- or 4tU-labeled RNA in the total RNA pool (7). Here, newly synthesized RNA is similarly labeled with 4sU or 4tU. Total RNA is purified and treated with the thiol-reactive alkylating reagent iodoacetamide, which chemically recodes labeled RNA and results in the misincorporation of guanosine during reverse transcription. Thus, thymine-to-cytosine (T>C) conversions define newly synthesized RNA and are quantified using dedicated data analysis packages (e.g., SLAM-DUNK or GRAND-SLAM) or alternative approaches (23–26). SLAM-seq has several advantages compared to 4tU-seq, including increased reproducibility as the purification of labeled RNA is not required. Additionally, as SLAM-seq captures both modified and unmodified RNA, reads derived from labeled RNA can be normalized by reads derived from total RNA, which can bypass the need for external spike-in normalization even when global changes in transcription are observed (8). SLAM-seq and similar methods have been used in *S. cerevisiae*, mainly to study the synthesis and decay of RNA (13, 23, 27).

Here, we provide an end-to-end workflow for rapidly degrading a target protein using the AID system and quantifying newly synthesized mRNA using SLAM-seq in *S. cerevisiae*. We include methods for targeted protein degradation, 4tU incorporation, rapid fixation, RNA purification, RNA alkylation, and RNA-seq library preparation. Although the individual protocols described here are not novel per se, this workflow provides a complete resource for turnkey implementation of these methods, which we believe will benefit others working with *S. cerevisiae*. In addition, this workflow is readily adaptable to other systems, including industrial or pathogenic fungi, and will benefit the larger research community.

## RESULTS AND DISCUSSION

### Engineering a workflow to study newly synthesized mRNA following rapid protein depletion in *Saccharomyces cerevisiae*

To address unresolved questions in transcription and other biological processes, we built an end-to-end workflow to rapidly deplete proteins of interest and measure newly transcribed mRNA. To support the implementation of this workflow, we provide an open-source, step-by-step protocol on protocols.io: DOI: dx.doi.org/10.17504/protocols.io.n2bvj3dj5lk5/v1

### Applying the SLAM-seq workflow to study the transcriptional regulators Bdf1 and Bdf2 in *Saccharomyces cerevisiae*

To test our workflow, we performed experiments with two *S. cerevisiae* strains (RDY73 and RDY234) related to our studies on BET proteins. BET (bromodomain and extra-terminal domain) proteins are conserved chromatin readers characterized by two bromodomains and an extra-terminal (ET) domain (28). BET proteins have an integral role in transcriptional regulation, although the mechanisms by which they regulate transcription are not well understood (9, 28–30). There are two BET proteins (Bdf1 and Bdf2, referred to here as Bdf1/2) in *S. cerevisiae*, and deletion of both *BDF1* and *BDF1* causes synthetic lethality. Although Bdf1 and Bdf2 have overlapping functions, the depletion of Bdf1 has a more significant impact on global transcription (9). Strain RDY73 carries AID-tagged BDF1/2 and was previously used to study changes in transcription after depletion of Bdf1/2 (9). To obtain strain RDY234, an additional, unmodified copy of *BDF1* driven by its endogenous promoter was integrated into the *TRP1* locus of strain RDY73.

In this study, cells of strain RDY73 or RDY234 in the logarithmic phase were treated with the auxin 3-indoleacetic acid (IAA) for 25 min to deplete Bdf1/2 then treated with thiouracil (4tU) for 4 min to label newly transcribed RNA. Immediately after 4tU labeling, cells were rapidly fixed in methanol on dry ice as described (22, 31, 32). Depletion of Bdf1/2 was validated by western blotting (data not shown). Subsequently, RNA was purified, alkylated, reverse transcribed, and sequenced. To assess how well our workflow quantifies 4tU-labeled transcripts, we first calculated percent conversion rates in reads mapped to defined windows. We observed a strong and specific accumulation of T>C conversions in 4tU-treated cells compared to untreated cells (Figure 1A). Second, we calculated percent read counts with 0, 1, or ≥ 2 T>C conversions mapped to defined windows. We observed that most reads have no T>C conversions in both 4tU-treated and untreated cells and that, although a percentage of reads with one T>C conversion in untreated cells was observed, 4tU-treated cells have a higher percentage of reads with one or ≥ 2 T>C conversions compared to untreated cells (Figure 1B-C). We also calculated the percent background signal in reads with ≥ 1 or 2 T>C conversions mapped to defined windows. Here, we observed a substantially lower percent background signal in reads with ≥ 2 T>C conversions compared to reads with ≥ 1 T>C conversions (Figure 2D). Together, these results illustrate that our workflow can selectively quantify 4tU-labeled transcripts and that counting reads with ≥ 2 T>C conversions adequately reduces background signal.

**Figure 1.**
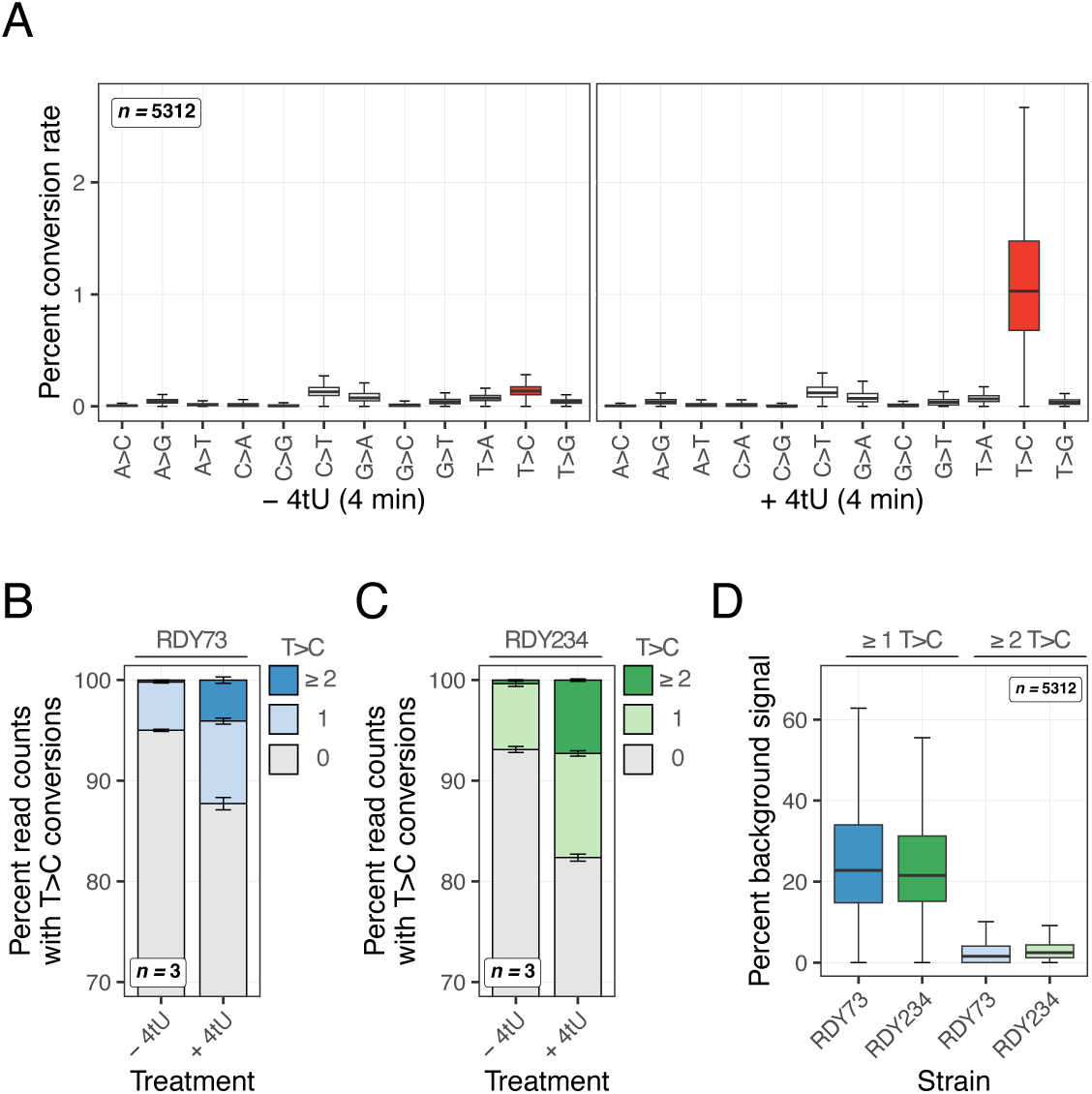
Application of SLAM-seq to *Saccharomyces cerevisiae*. (A) Percent conversion rates in reads mapped to defined windows. SLAM-seq data was derived from *S. cerevisiae* cells (strain RDY73) treated with (+) or without (−) thiouracil (4tU) for 4 min. Conversion rates across the indicated number of transcripts (*n =* 5312) are presented as Tukey boxplots. Outliers are not shown. (B) Percent read counts with 0, 1, or ≥ 2 T>C conversions mapped to defined windows. SLAM-seq data was derived from *S. cerevisiae* cells (strain RDY73) treated with (+) or without (−) thiouracil (4tU) for 4 min. Error bars represent the standard deviation of the mean (*n =* 3). The y-axis begins at 70 percent. (C) Percent read counts with 0, 1, or ≥ 2 T>C conversions mapped to defined windows. SLAM-seq data was derived from *S. cerevisiae* cells (strain RDY234) treated with (+) or without (−) thiouracil (4tU) for 4 min. Error bars represent the standard deviation of the mean (*n =* 3). The y-axis begins at 70 percent. (D) Percent background signal in reads with ≥ 1 or 2 T>C conversions mapped to defined windows. SLAM-seq data was derived from *S. cerevisiae* cells (strains RDY73 or RDY234) treated with or without 4tU for 4 min. Background signals across the indicated number of transcripts (*n =* 5312) are presented as Tukey boxplots. Outliers are not shown.

**Figure 2.**
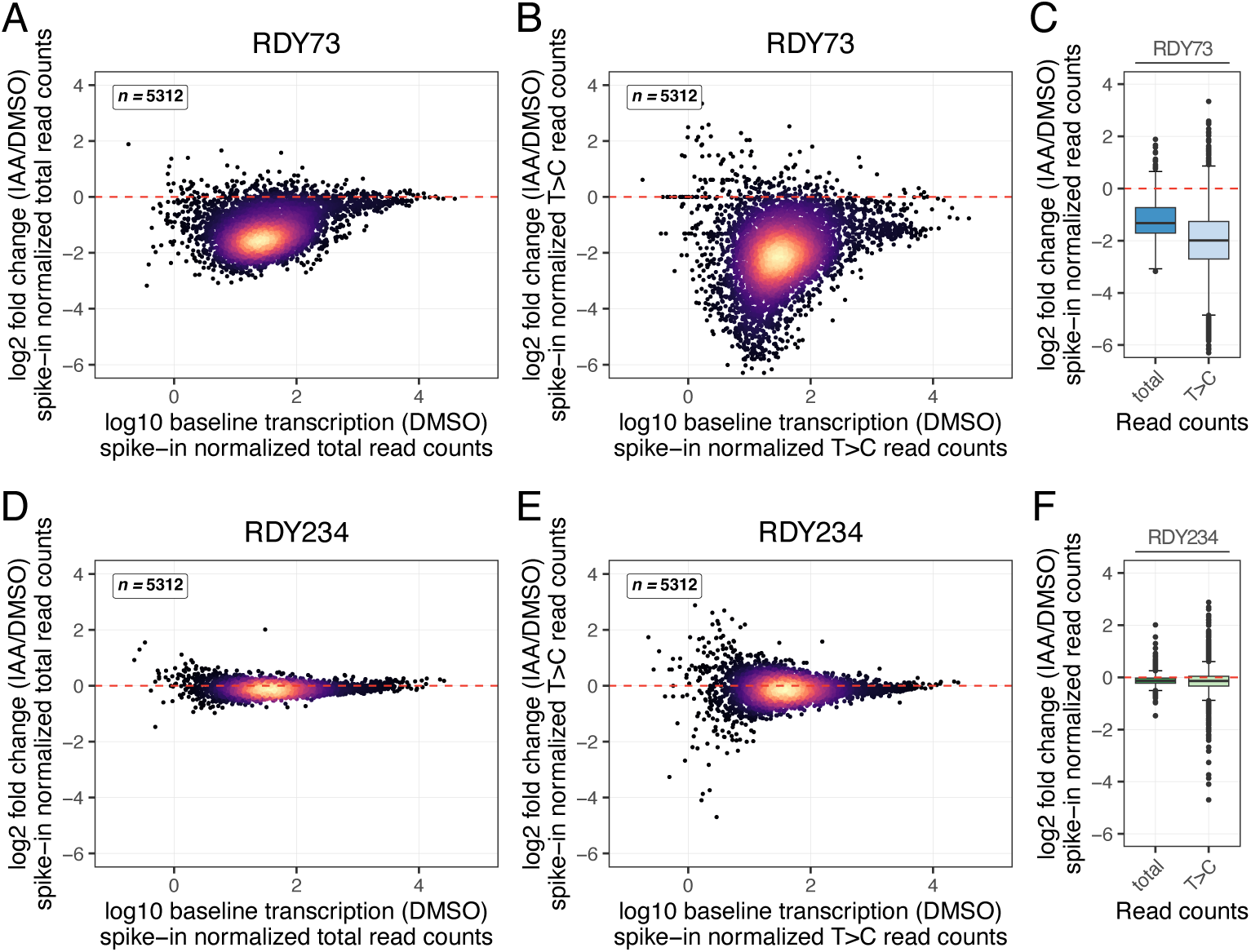
SLAM-seq detects global transcriptional changes associated with rapid depletion of Bdf1/2. (A) Scatterplot comparing log2 fold change in transcription and log10 baseline transcription following Bdf1/2 depletion (strain RDY73) across the indicated number of transcripts (*n =* 5312). Total read counts were normalized using size factors calculated on total read counts from exogenous whole cell spike-in. (B) Scatterplot comparing log2 fold change in transcription and log10 baseline transcription following Bdf1/2 depletion (strain RDY73) across the indicated number of transcripts (*n =* 5312). T>C read counts were normalized using size factors calculated on total read counts from exogenous whole cell spike-in. (C) Boxplots comparing log2 fold changes in transcription following Bdf1/2 depletion (strain RDY73) normalized using exogenous whole cell spike-in. Changes in transcription across 5312 transcripts presented as Tukey boxplots. (D) Scatterplot comparing log2 fold change in transcription and log10 baseline transcription following Bdf1/2 depletion in a strain ectopically expressing *BDF1* (RDY234) across the indicated number of transcripts (*n =* 5312). Total read counts were normalized using size factors calculated on total read counts from exogenous whole cell spike-in. (E) Scatterplot comparing log2 fold change in transcription and log10 baseline transcription following Bdf1/2 depletion in a strain ectopically expressing *BDF1* (RDY234) across the indicated number of transcripts (*n =* 5312). T>C read counts were normalized using size factors calculated on total read counts from exogenous whole cell spike-in. (F) Boxplots comparing log2 fold changes in transcription following Bdf1/2 depletion in a strain ectopically expressing *BDF1* (RDY234) normalized using exogenous whole cell spike-in. Changes in transcription across 5312 transcripts presented as Tukey boxplots.

### SLAM-seq is robust to normalization using whole-cell spike-in or total read counts

#### Normalization by whole-cell spike-in normalization

In initial SLAM-seq experiments, we performed whole-cell spike-in normalization using 4tU-labeled *S. pombe* cells as previously described (9, 12). Here, we calculated spike-in normalization factors for each sample as the sum of all *S. pombe* reads mapped divided by 100,000. The factor 100,000 was arbitrarily chosen to ensure the spike-in normalization factors for most samples were between one and ten. We then used the spike-in normalization factors to normalize (1) total read counts or (2) T>C read counts in DESeq2. Interestingly, we observed a global loss of transcription following Bdf1/2 depletion for both spike-in normalized total read counts and spike-in normalized T>C read counts derived from strain RDY73 (Figure 2A-B). These results indicate that substantial changes in transcription are apparent in steady-state RNA 30 min after inducing Bdf1/2 depletion. However, the magnitude of the transcriptional changes is much more dramatic in newly transcribed RNA compared to steady-state RNA (Figure 2C). We did not observe global changes in transcription for either spike-in normalized total read counts or spike-in normalized T>C read counts derived from strain RDY234 (Figure 2D-F), indicating that ectopically expressed *BDF1* largely complements depletion of endogenous Bdf1/2. Taken together, these results illustrate the importance of measuring newly transcribed RNA to study the direct effects of a perturbation such as rapid protein depletion.

We additionally tested normalization using *S. pombe* reads with a minimum of 2 T>C conversions rather than total *S. pombe* reads, though no substantial benefit was found using this approach (data not shown). Thus, we propose that SLAM-seq largely circumvents the need to use 4tU-labeled spike-in. However, as a general consideration, we note that 4tU-labeled spike-in is needed if samples will also be used for 4tU-seq or TT-SLAM-seq (33).

#### Normalization by reads derived from total RNA

In previous studies, SLAM-seq data has been normalized using reads derived from total RNA rather than whole-cell spike-in (8, 34). These studies have demonstrated that normalizing T>C read count data by total read count data is a robust approach even when global changes in gene transcription are observed. In addition, potential issues with spike-in normalization have been noted (35, 36). Therefore, we wanted to test a normalization approach using reads derived from total RNA to circumvent the need for whole-cell spike-in and streamline our workflow.

Previous studies used DESeq2 to calculate size factors based on total read counts. The DESeq2 size factors were then used to normalize T>C read counts. We initially applied this approach to our RDY73 data. We observed a global loss of transcription following Bdf1/2 depletion for DESeq2 normalized T>C read counts (Figure 3A). However, the magnitude of these changes appeared to be less than what we observed for spike-in normalized T>C read counts. The median-ratio method implemented by DESeq2 to estimate size factors assumes that the expression of most genes is not affected by experimental conditions (37). We presume that the extent to which Bdf1/2 depletion affects steady-state transcription in our experiments may violate this assumption.

**Figure 3.**
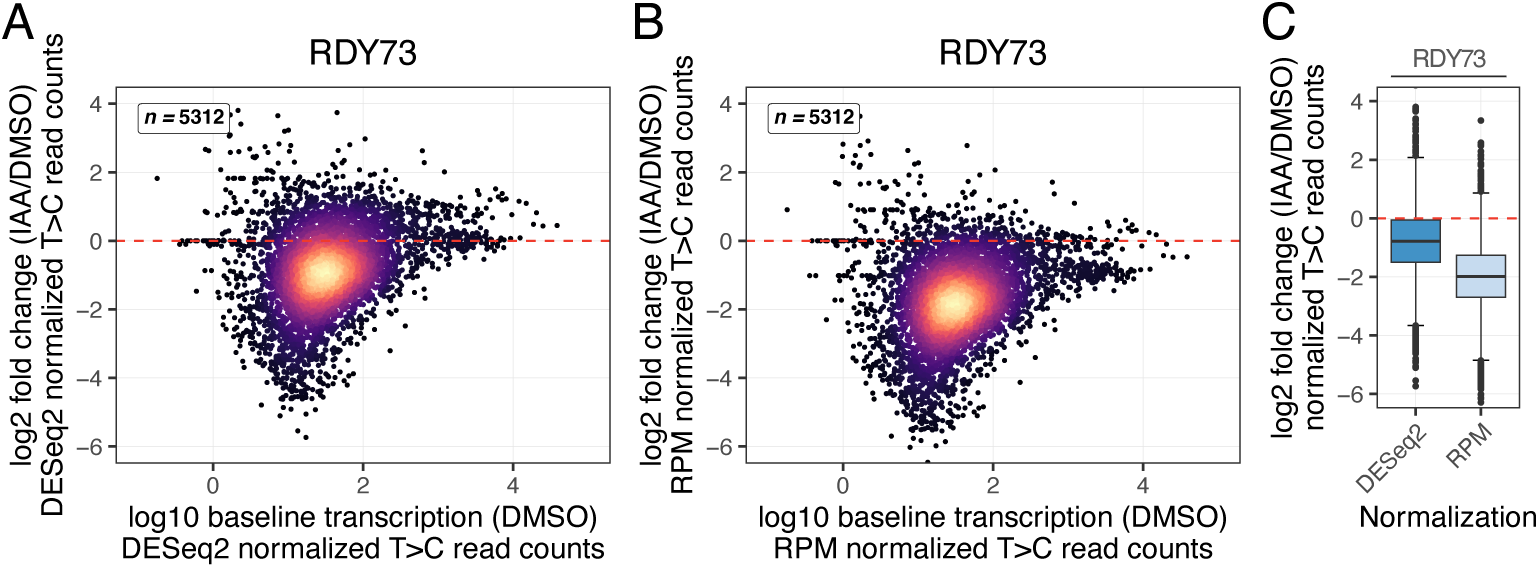
SLAM-seq is robust to normalization using total read counts. (A) Scatterplot comparing log2 fold change in transcription and log10 baseline transcription following Bdf1/2 depletion across the indicated number of transcripts (*n =* 5312). T>C read counts were normalized using size factors estimated on total read counts using DESeq2. (B) Scatterplot comparing log2 fold change in transcription and log10 baseline transcription following Bdf1/2 depletion across the indicated number of transcripts (*n =* 5312). T>C read counts were normalized using reads per million (RPM) size factors calculated on total read counts. (C) Boxplots comparing log2 fold changes in transcription following Bdf1/2 depletion normalized using total read counts. Changes in transcription across 5312 transcripts are presented as Tukey boxplots.

Based on these results, we reasoned that an alternative normalization approach using total read count data might yield better results for a rapidly dividing organism like *S. cerevisiae*. Thus, we calculated reads per million (RPM) normalization factors based on total read counts for each sample as the sum of all reads mapped divided by one million. We then used these RPM normalization factors to normalize T>C read counts in DESeq2. Using this approach, we again observed a global loss of transcription following Bdf1/2 depletion (Figure 3B). However, the magnitude of transcriptional changes was (1) more dramatic when T>C read counts were normalized by RPM compared to DESeq2 size factors (Figure 3C) but (2) very similar when T>C read counts were normalized by RPM compared to DESeq2 size factors. Together, these results illustrate that SLAM-seq data derived from *S. cerevisiae* can be reliably normalized using RPM normalization factors derived from total read counts.

### SLAM-seq correlates well with 4tU-seq and identifies extensive differential expression following depletion of Bdf1 and Bdf2

Next, we compared our SLAM-seq data to published 4tU-seq data derived from strain RDY73 under comparable experimental conditions (9). A striking loss of transcription is apparent in the data from both SLAM-seq and 4tU-seq (Figure 3B) (9). Comparing the change in transcription across the two datasets, we observe a strong correlation in SLAM-seq and 4tU-seq experiments. Furthermore, we observe extensive differential expression following Bdf1/2 depletion (Figure 4B). For example, we detected nine upregulated genes and 2720 downregulated genes using a false discovery rate (FDR) adjusted p-value ≤ 0.05 and log2 fold change ≥ |1| as cutoffs. Together, these results illustrate a strong correlation between the results of SLAM-seq and 4tU-seq and demonstrate that our workflow can help define the direct targets of transcriptional regulators such as Bdf1/2 in *S. cerevisiae*.

**Figure 4.**
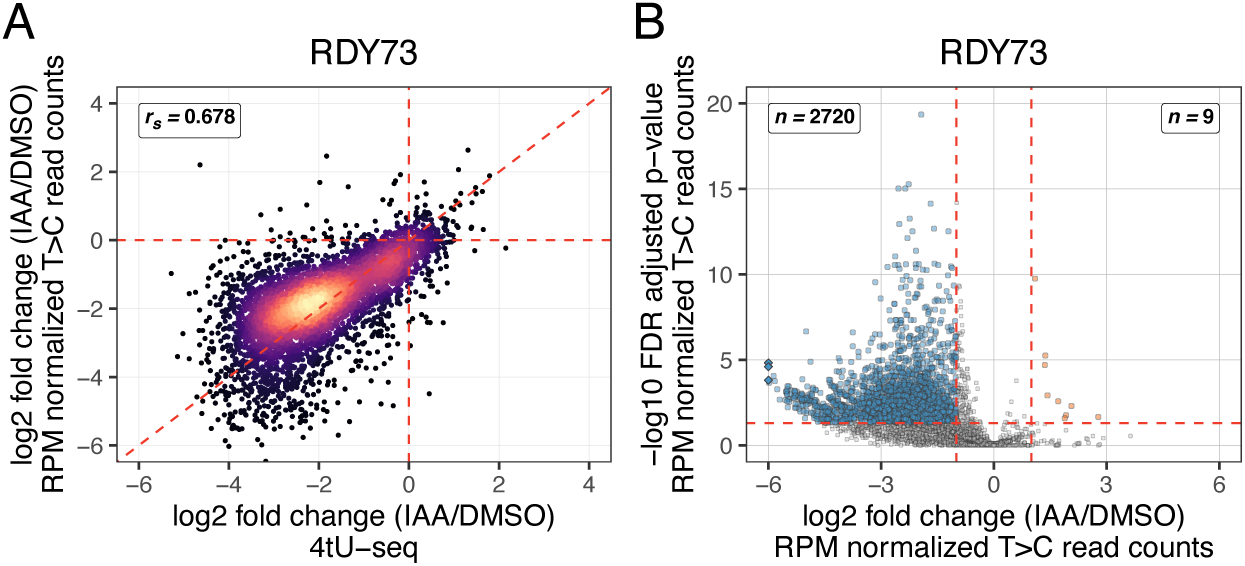
SLAM-seq correlates well with 4tU-seq and identifies extensive differential expression following Bdf1/2 depletion. (A) Scatterplot comparing log2 fold change in transcription following Bdf1/2 depletion determined using SLAM-seq or 4tU-seq across 4883 transcripts. T>C read counts derived from SLAM-seq were normalized using reads per million (RPM) size factors calculated on total read counts. Read counts derived from previously published 4tU-seq data were normalized using size factors calculated on total read counts from exogenous whole cell spike-in. Spearman’s correlation coefficient (*r*_s_) is shown. (B) Volcano plot comparing −log10 false discovery rate (FDR) adjusted p-value and log2 fold change in transcription following Bdf1/2 depletion across 5312 transcripts. T>C read counts were normalized using reads per million (RPM) size factors calculated on total read counts. Numbers of differentially expressed genes are shown in respective regions of the plot.

### Limitations

First, we and others (8) provide data that a conversion per read threshold of ≥ 2 T>C conversions adequately reduces background signal. However, we recommend that initial SLAM-seq experiments include an untreated (–4tU) control to assess spurious nucleotide conversions and polymorphisms between experimental and reference strains. Second, methods for preparing mRNA-seq libraries that use a template switching oligo (TSO) such as that described here generate forward reads that start with three low-diversity bases. It is therefore necessary to increase the diversity of the sequencing pool. Final sequencing libraries should be pooled with 10-15% libraries prepared using standard approaches or control libraries (see (38) for additional details). Furthermore, similar to other 3′ mRNA-seq methods (e.g., QuantSeq; Lexogen, Greenland, NH, USA) the reverse read provides minimal useful information. The method described here uses barcode information in both the forward and reverse reads. In our experience, paired-end sequencing can be more cost-effective than single-end sequencing depending on the provider. Third, while we anticipate this workflow will be readily adapted to other systems, we acknowledge that some optimization will be necessary. Using parts of the workflow described here, we have successfully prepared 3′ mRNA-seq libraries from filamentous fungi and human cell lines (data not shown).

### Conclusions

This work provides an end-to-end workflow for rapid and specific protein degradation using the AID system and selective quantification of newly synthesized RNA using SLAM-seq in *S. cerevisiae*. The workflow builds on previous methods to provide a complete resource for implementation. We demonstrate that the workflow can help define the direct effects of transcriptional regulators using the BET proteins Bdf1 and Bdf2 as a test case. As such, this workflow will help address outstanding questions underlying the molecular basis of transcription and other biological processes in *S. cerevisiae*. Additionally, this workflow is adaptable to other systems, including industrial, pathogenic, or other model fungi, which will benefit the larger research community.

## MATERIALS AND METHODS

### Strains and general culturing conditions

All *S. cerevisiae* and *S. pombe* strains used in this study are listed in Supplementary Table 2. Strain RDY234 was engineered using standard methods. *Saccharomyces cerevisiae* was grown in YPD medium (10 g/l yeast extract, 20 g/l peptone, 20 g/l glucose, 20 mg/ml adenine sulfate) at 30°C with shaking at 220 rpm. *Schizosaccharomyces pombe* was grown in YE medium (5 g/l yeast extract, 30 g/l glucose) at 30°C with shaking at 220 rpm. *Saccharomyces cerevisiae* was grown to a final OD600 of ∼0.7 and *S. pombe* was grown to a final OD600 of ∼1.0. Three biological replicates were collected in all experiments.

### IAA treatment, 4tU labeling, sampling, and rapid fixation

*Saccharomyces cerevisiae* was grown in 40 ml of YPD at 30°C with shaking at 220 rpm. The 40 ml cultures were grown to an OD600 of ∼0.45 and split into three 10 ml cultures. Cultures were equilibrated for 5 min at 30°C with shaking at 220 rpm. Following equilibration, 50 μl of 0.2 M 3-indoleacetic acid (IAA) freshly prepared in dimethyl sulfoxide (DMSO) or DMSO was added to the appropriate culture; cultures were immediately and vigorously mixed and incubated for 25 min at 30°C with shaking at 220 rpm. Immediately following incubation, 25 μl of 2 M 4-thiouracil (4tU) freshly prepared in dimethyl sulfoxide (DMSO) or DMSO was added to the appropriate culture; cultures were immediately and vigorously mixed and incubated for exactly 4 min at 30°C with shaking at 220 rpm. Immediately after 4tU treatment, the cultures were decanted into 5 ml of 100% methanol in a 50 conical tube prechilled on dry ice. The slurry was gently mixed by swirling to ensure homogenization and kept on dry ice. An aliquot of the slurry was collected for cell counting and western blotting. The remaining slurry was centrifuged at 3,000 g and 4°C for 10 min to pellet the cells. The supernatant was decanted, and cells were resuspended in 400 μl of DNA/RNA shield (Zymo Research, Irvine, CA, USA) by pipetting. Cell suspensions were flash frozen on dry ice and stored at –80°C.

### RNA purification and DNase I treatment

RNA purification and on-column DNase I treatment were performed using a Quick-RNA Fungal/Bacterial Miniprep kit and DNase I set (Zymo Research) following the manufacturer’s recommendations with minor modifications. Samples were protected from light when possible (i.e., samples were processed under low light and/or covered with foil). Reagents (i.e., RNA lysis buffer, RNA wash buffer, RNA prep buffer, 100% ethanol, and nuclease-free water) were supplemented with dithiothreitol (DTT; GoldBio, Saint Louis, MO, USA) to maintain reducing conditions. DNA digestion buffer was not supplemented with DTT. Samples stored in DNA/RNA shield at –80°C were thawed at room temperature, and an equal volume of RNA lysis buffer (supplemented with DTT) was added to each sample. Samples were homogenized twice in a Mini-Beadbeater-24 (Biospec Products, Bartlesville, OK, USA) at 3800 rpm for 45 s immediately followed by incubation on ice for 2 min. DNase I-treated total RNA was eluted in 50 μl of nuclease-free water (supplemented with DTT) prewarmed to 50°C.

### RNA alkylation

RNA alkylation was performed as previously described (7, 13) with minor modifications. Samples were protected from light when possible (i.e., samples were processed under low light and/or covered with foil). A total of 5 μg of total RNA in 20 μl of nuclease-free water was used for alkylation. The RNA was combined with 5 μl of 100 mM iodoacetamide (Sigma-Aldrich, St. Louis, MO, USA), 5 μl of 0.5 M sodium phosphate buffer (pH 8.0), and 20 μl of DMSO. The reaction was gently mixed and incubated in a thermomixer at 900 rpm and 50°C for 15 min in the dark. A volume of 1 μl of 1 M DTT was added to stop the reaction. Exposure to light was considered acceptable following the addition of DTT. Alkylated RNA was precipitated as previously described (7) and resuspended in 30 μl of nuclease-free water.

### 3′ mRNA-sequencing library preparation

3′ mRNA-sequencing library preparation is based on previously described methods with modifications (38, 39). A total of 200 ng of alkylated RNA in 5 μl of nuclease-free water was used for library preparation. The remaining steps of library preparation are provided in detail in the full step-by-step protocol (https://www.protocols.io/). Libraries were sequenced on a NovaSeq 6000 (Illumina, San Diego, CA, USA) at the Oklahoma Medical Research Foundation (OMRF) Clinical Genomics Center (Oklahoma City, OK, USA).

### Data analysis

Demultiplexed, paired-end 150 bp reads were preprocessed using fastp (version 0.23.2) (40) to extract unique molecular identifiers (UMIs) and BBMap (version 36.99) (https://sourceforge.net/projects/bbmap/) to trim adapter sequences and polyA tails. The reverse reads generated from most 3′ mRNA-seq methods provide minimal information beyond barcoding. Therefore, only forward reads were then processed using SLAM-DUNK (version 0.4.3) (25). Within SLAM-DUNK, reads were aligned using NextGenMap (version 0.5.5) (41) and single-nucleotide polymorphisms (SNPs) were called using VarScan 2 (version 2.4.5) (42). To define counting windows for SLAM-DUNK, a BED file was created containing all annotated open reading frames (ORFs) in the *S. cerevisiae* strain S288C assembly (version R64-3-1) using BEDOPS (version 2.4.3) (43). All ORFs were extended 250 bp beyond their stop position to capture 3′ untranslated regions (UTRs) using BEDTools (version 2.30.0) (44). Total read counts are defined as all reads remaining after alignment filtering and recovery of multimapping reads in SLAM-DUNK. Unless otherwise indicated, a conversion per read threshold of ≥ 2 T>C conversions was used to define T>C read counts.

Genes classified as dubious, pseudogenes, or transposable elements were excluded from subsequent analyses. The remaining genes were filtered against 5313 genes that were previously found to be reliably detected under experimental conditions comparable to those used in this study (9), which left 5312 genes. To compare SLAM-seq and 4tU-seq data, genes were further filtered against 4883 genes that were previously found to provide high reproducibility and low information loss in 4tU-seq experiments with strain RDY73 (9), which left 4883 genes. Differential gene expression analysis was performed on total read counts and T>C read counts using DESeq2 (version 1.38.3) (37). Adjusted p-values were calculated using IHW (version 1.26.0) (45) within DESeq2. Data processing and plotting were performed using R (version 4.2.3) (https://www.R-project.org/) in RStudio (version 2023.12.0+369) (http://www.rstudio.com/).

### Data availability

Sequencing data generated in this study were deposited in the NCBI Gene Expression Omnibus (GEO) repository as GEO Dataset GSEXXXXXX and are additionally accessible from the NCBI Sequence Read Archive (SRA) under BioProject PRJNAXXXXXX. The reference genome assembly and annotation for *Saccharomyces cerevisiae* strain S288C (version R64-3-1, RefSeq Assembly GCF_000146045.2) were retrieved from the NCBI Datasets repository. Published sequencing data (GEO Dataset GSE171067) were retrieved from the NCBI GEO repository. All other relevant data supporting key findings in this study are available within the article or supplementary information.

## ACKNOWLEDGMENTS

We thank Magdalena Donczew for assistance in generating genetic resources and Magdalena Donczew and Agnieszka Machowska for helpful discussion and critically reviewing the manuscript. We thank the OMRF Clinical Genomics Center for helpful discussion and high-throughput sequencing and the OMRF Center for Biomedical Data Sciences for helpful discussion.

## SUPLEMENTARY MATERIAL

**Supplementary Table 1.** Summary of sequencing data generated in this study.

**Supplementary Table 2.** *Saccharomyces cerevisiae* and *Schizosaccharomyces pombe* strains used in this study.

